# Clinically adaptable polymer enables simultaneous spatial analysis of colonic tissues and biofilms

**DOI:** 10.1101/2020.04.15.030874

**Authors:** Mary C. Macedonia, Julia L. Drewes, Nicholas O. Markham, Alan J. Simmons, Joseph T. Roland, Paige N. Vega, Cherie’ R. Scurrah, Robert J. Coffey, Martha J. Shrubsole, Cynthia L. Sears, Ken S. Lau

**Affiliations:** Department of Medicine, Division of Gastroenterology, Hepatology and Nutrition, Vanderbilt University Medical Center, Nashville TN; Epithelial Biology Center, Vanderbilt University Medical Center, Nashville TN; Department of Medicine, Division of Infectious Diseases, Johns Hopkins University School of Medicine, Baltimore MD; Department of Cell and Developmental Biology, Vanderbilt University School of Medicine, Nashville TN; Department of Surgery, Vanderbilt University Medical Center, Nashville TN; Vanderbilt Ingram Cancer Center, Nashville, TN; Department of Medicine, Division of Epidemiology, Vanderbilt Epidemiology Center, Vanderbilt University Medical Center, Nashville TN

## Abstract

Microbial influences on host cells depend upon the identities of the microbes, their spatial localization, and the responses they invoke on specific host cell populations. Multi-modal analyses of both microbes and host cells in a spatially-resolved fashion would enable studies into these complex interactions in native tissue environments, potentially in clinical specimens. While techniques to preserve each of the microbial and host cell compartments have been used to examine tissues and microbes separately, we endeavored to develop approaches to simultaneously analyze both compartments. Herein, we established an original method for mucus preservation using Poloxamer 407 (also known as Pluronic F-127), a thermoreversible polymer with mucus-adhesive characteristics. We demonstrate that this approach can preserve spatially-defined compartments of the mucus bi-layer in the colon and the bacterial communities within, compared with their marked absence when tissues were processed with traditional formalin-fixed paraffin-embedded (FFPE) pipelines. Additionally, antigens for antibody staining of host cells were preserved and signal intensity for 16S rRNA fluorescence *in situ* hybridization (FISH) was enhanced in Poloxamer-fixed samples. This in turn enabled us to integrate multi-modal analysis using a modified multiplex immunofluorescence (MxIF) protocol. Importantly, we have formulated Poloxamer 407 to polymerize and crosslink at room temperature for use in clinical workflows. These results suggest that the fixative formulation of Poloxamer 407 can be integrated into biospecimen collection pipelines for simultaneous analysis of microbes and host cells.

## Introduction

Investigating host-microbe interactions may reveal the mechanisms underlying how changes in the gastrointestinal microbiome are related to colorectal cancer (CRC). Recently, increased abundance of *Fusobacterium nucleatum, Bacteroides fragilis*, and *Escherichia coli* have been associated with human CRC ^1-3^. These species are hypothesized to promote tumorigenesis and/or progression of CRC through up-regulation of oncogenes, modification of intestinal mucus, or damage to host DNA^2,4,5^. In addition to the mere presence or absence of putative carcinogenic bacteria in the gut, the spatial localization and physical interaction with host epithelial cells may be critically important.

Bacterial biofilms are ubiquitous in nature and can also be observed routinely in the gastrointestinal tract of healthy individuals specifically in the outer layer of the MUC2-dominated mucus bi-layer that lines the luminal colonic surface^6-11^. In contrast, the inner, striated mucus layer of the colon is largely sterile in healthy hosts ^6^. Perturbations of this tightly regulated spatial organization have been linked to both inflammatory bowel diseases (IBD) and CRC ^12-15^. Mucosa-associated bacterial biofilms are defined as microbial aggregates embedded within a polymeric matrix of both host and bacterial origin that directly contacts epithelial cells ^14,16^. Mucosa-associated biofilms have been identified in approximately 50% of sporadic CRC patients, 100% of familial adenomatous polyposis (FAP) patients, and 13% of healthy individuals. They are particularly enriched in CRCs from the ascending colon ^3^. Additionally, bacterial isolates from CRC or FAP biofilms have been shown to promote colonic tumor progression in genetically susceptible mice ^13,15^.

While many studies have characterized the microbiome through sequencing and/or mass spectrometry, host-microbe spatial relationships can only be studied with sufficient resolution through microscopy-based techniques, such as FISH or electron microscopy ^11,12^. Moreover, while sophisticated multimodal analyses are being conducted on the microbiome ^17^, concurrent analyses of host cells have lagged. Studies on the host response to microbiome alterations to date have typically relied on macroscopic outputs, such as tumor formation, or molecular assays performed on separate, spatially disconnected specimens ^2,15^. We propose that concurrent, multi-modal analysis of both host and microbial cells within the same tissue specimen would be a significant advance for deriving insights into host-microbe interactions in CRC, as well as other forms of pathogenesis.

Recently developed imaging techniques enable multiplexed *in situ* analysis of 50 or more analytes on the same tissue section ^18-27^. However, one challenge has been identifying a tissue processing method that enables simultaneous spatial localization and quantifiable detection of both host components, such as proteins, and bacterial components via immunofluorescence (IF) and FISH, respectively. The colonic mucus layer acts as both a resident niche and a barrier for the luminal microbiome, playing well-documented roles in colonic pathologies such as ulcerative colitis ^10,14,28,29^. The hydrophilic colonic mucus is easily disrupted by hydration and dehydration of formalin-fixed tissue with standard histological preparations ^7,30^. Carnoy’s solution is a rapidly dehydrating fixative that preserves mucus ^9,14^, but it reduces the sensitivity of IF and FISH ^7,31^. Our goal for this work is to devise a tissue processing strategy that preserves mucus architecture and is amenable to coincident antibody and nucleic acid detection. Herein, we describe how Poloxamer 407 fixation permits the simultaneous characterization of host cell protein expression and identification of microbial communities within the adjacent colonic mucus. Furthermore, we deployed this approach into a pilot clinical pipeline to enable multiplex *in situ* profiling of human tissues and associated microbes.

## Results

### Carnoy’s and formalin fixation are incompatible with simultaneous mucus preservation and host cell assays

The alcohol-based Carnoy’s solution (Carnoy’s fixative/Methacarn) is a rapid dehydration fixative that preserves the hydrophilic mucus layer for colonic microbial FISH analysis. Cross-linking fixative, such as neutral buffered formalin (NBF), is used for standard FFPE tissue processing and has optimal antigen preservation for immunofluorescence (IF) or immunohistochemistry (IHC) ^32,33^. For this study, we aimed to identify the best tissue fixation method for simultaneous microbial FISH and host cell IF. We initially used mouse colon tissue as a model because of its abundance and availability. Mice do not naturally form mucosa-associated biofilms, but their mucus layers are arranged in bi-layer structures with different microbial compositions that can easily be evaluated.

Initially, we compared Carnoy’s fixation and NBF fixation, followed by standard histological processing. Standard IF stains such as proliferating cell nuclear antigen (PCNA) and the epithelial marker pan-cadherin (PCAD) had lower fluorescent signals in Carnoy’s-fixed mouse colon compared with NBF fixation of the same tissue (Fig. 1A-B). In contrast, NBF fixation results in loss of the mucopolysaccharide-containing mucus layer (Fig. 1C), because it cannot withstand hydration and shear forces during processing. NBF contains mostly water and solubilizes the hydrophilic mucus layer during the fixation process^11,34,35^. Consistent with known knowledge, these conventional fixatives are inadequate for *in situ* analysis of mucus-residing bacteria and host cells within the same tissue section.

**Figure 1:**
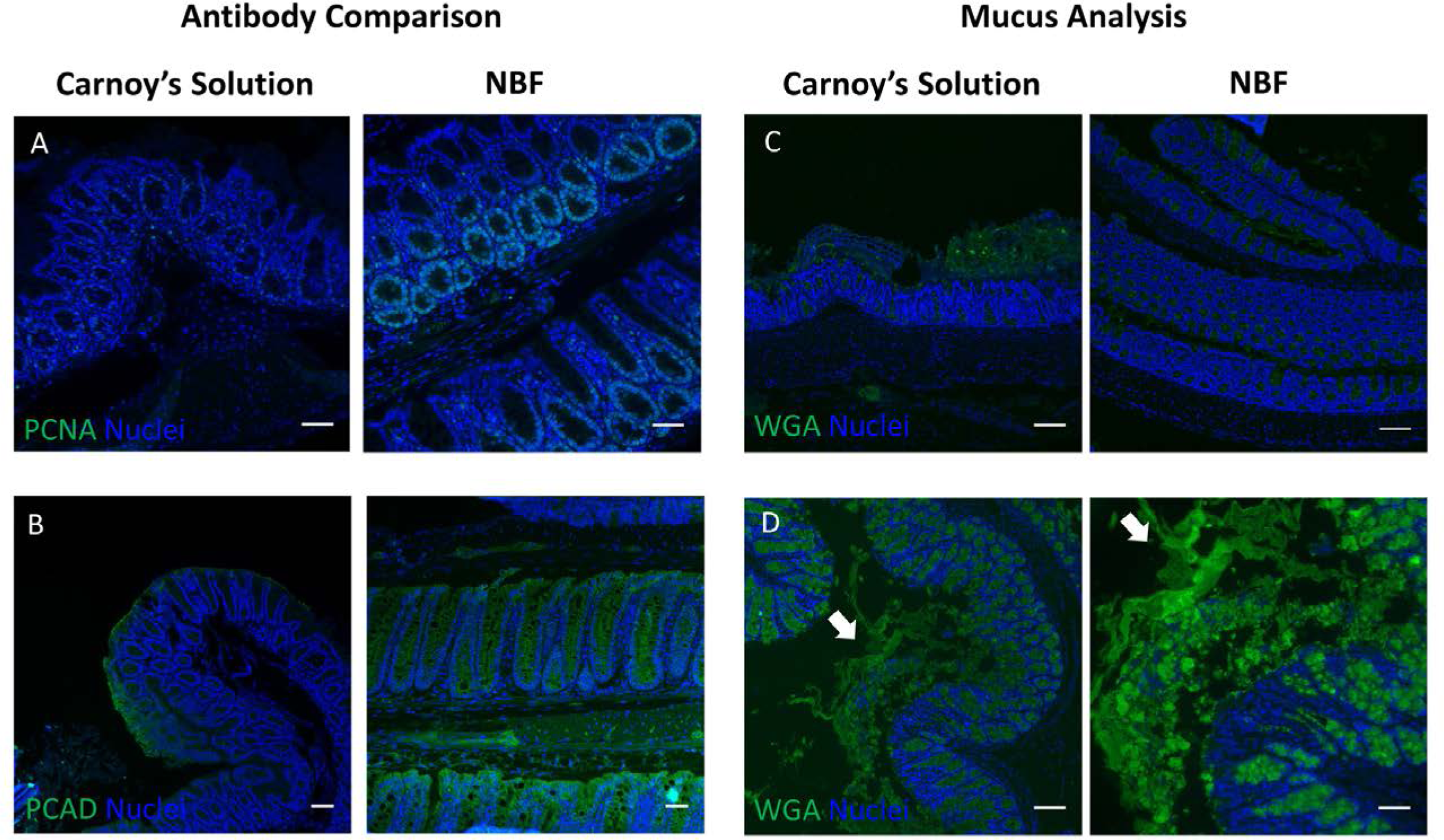
Carnoy’s and formalin fixation preclude multi-modal analysis of colonic tissues with intact mucus. Comparison of protein staining using IF between mouse colonic tissues fixed with Carnoy’s solution (left) or NBF (right) for (A) PCNA and (B) pan-Cadherin (PCAD). Scale bars = 50 μm (C) Mucus layer preservation as assessed by lectin (WGA) IF of mouse colonic tissues processed with Carnoy’s solution (left) or NBF (right). Scale bars = 100 μm (D) Mucus layer preservation using low melting point agarose as a scaffold. White arrows point to distorted portions of the mucus layer. Left - low magnification view. Scale bar = 100 μm. Left - high magnification view. Scale bar = 50 μm.

Scaffolding agents have been used to maintain the mucus layer during formaldehyde-based fixation^36^. Thus, we embedded freshly harvested mouse colon with low melting point agarose gel to scaffold the mucus while fixing the tissue with 4% paraformaldehyde at room temperature (Fig. 1D). The mucus layer was retained, consistent with previous studies, but its morphology was distorted by hydration. This modest improvement prompted us to identify substrates for scaffolding the mucus layer, including those that can chemically adhere and cross-link mucus.

### Poloxamer 407 formulated as a fixative solution polymerizes at room temperature

Poloxamer 407 is a bio-adhesive polymer with an A-B-A block configuration, with a hydrophobic polypropylene oxide (PPO) “B” block flanked by hydrophilic polyethylene oxide (PEO) “A” blocks (Fig. 2 A,B). Poloxamer 407 exhibits temperature-dependent and reversible polymerization essentially transforming from a viscous liquid at 4 C to a solid gel at higher temperatures capable of scaffolding biological surfaces. As temperature increases, hydrophobic B blocks align at the core while hydrophilic A blocks are oriented outward in a radial pattern to form micelles that can been used for nanoparticle drug delivery^37-40^. At the ceiling temperature, Poloxamer 407 forms a matrix of micelles held together by hydrogen bonds (Fig. 2C)^37,41^, with the outwardly arranged hydrogen molecules capable of forming non-covalent bonds with the highly glycosylated mucus matrix (Fig. 2D)^39^. Because of this transition into a gel state capable of providing scaffold support, we surmised that it would support the hydrophilic mucus architecture in histological applications.

**Figure 2:**
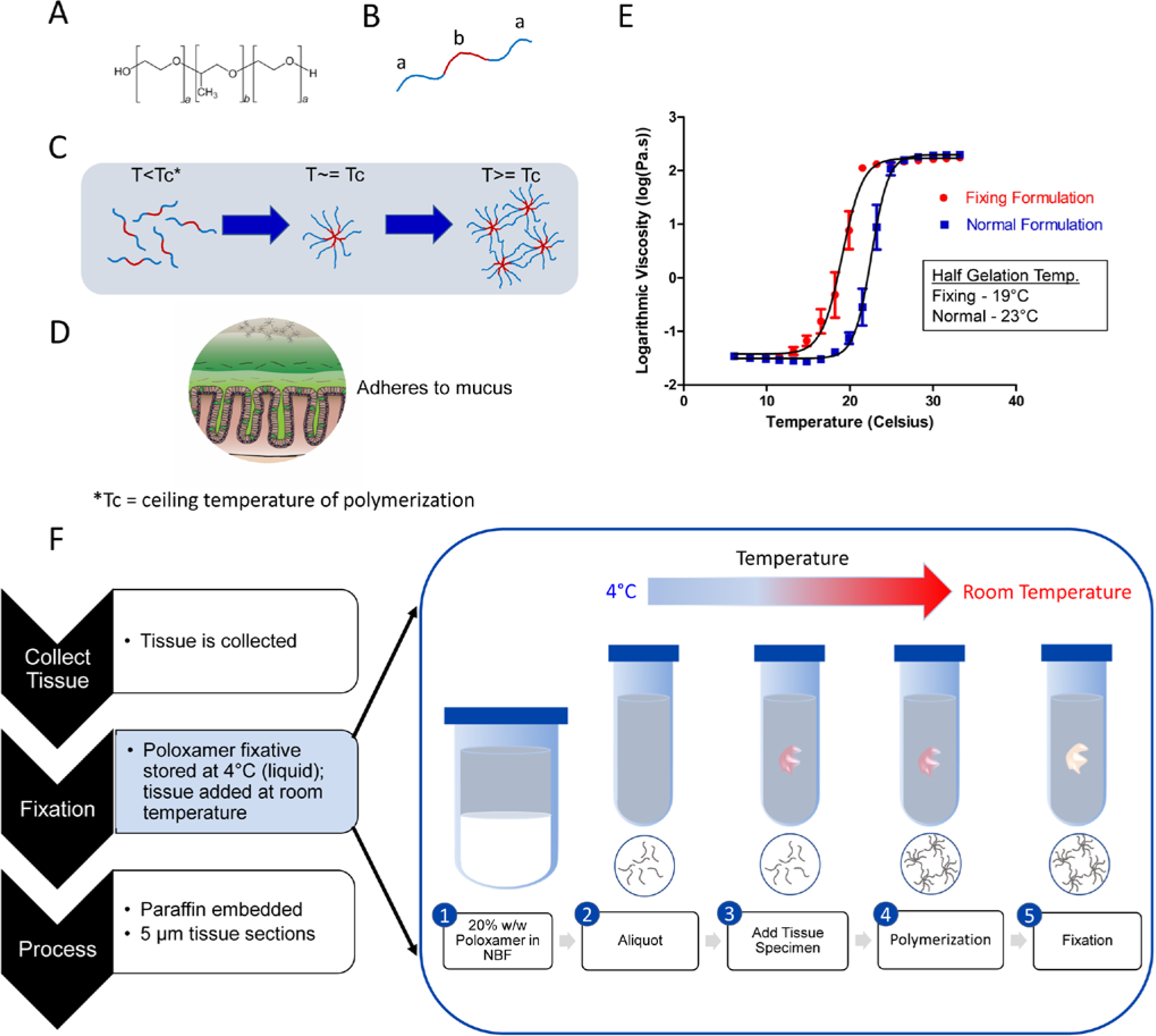
Formulation of Poloxamer 407 fixative polymerizes at room temperature. The structural characteristics (A) of Poloxamer 407 (or Pluronic F-127) as an A-B-A block thermoreversible polymer (B) with the “A” blocks of this polymer being hydrophilic (blue) and the “B” block being hydrophobic (red). (C) Temperature increases to the ceiling temperature (Tc) enable oriented micelle formation with b block interactions towards the center of the micelle, while at temperatures beyond the Tc, hydrogen bonds between different micelles facilitate the formation of a polymer matrix. (D) The polymer matrix has similar physical properties and also forms non-covalent bonds with glyco-moieties of the highly glycosylated mucosal matrix, allowing for efficient integration and scaffolding of Poloxamer 407 with the mucus layer. (E) Rheological testing to assess the polymer property of normal Poloxamer 407 (blue) and Poloxamer fixative (red) formulations as a function of temperature. Temperatures by which gelation is half maximum indicate a shift to a lower polymerization temperature comparing fixative versus normal formulations. (F) Histological workflow of tissue preparation with Poloxamer fixation. The fixation step in step 2 (e.g., with NBF) can be replaced with the Poloxamer fixative, while keeping the Poloxamer fixative on ice prior to addition to the tissue specimen. Fixation then undergoes the standard 24-hour fixation period at room temperature (right).

For its adaptation into the clinical workflow, we formulated the Poloxamer 407 solution such that it had 1) fixation properties in addition to structural support; and 2) a lower ceiling temperature such that it can be used at room temperature in clinical suites. These properties enable scaffolded fixation of a specimen in a single step without additional equipment or processing steps. We formulated Poloxamer 407 to 20% (w/w) in NBF solution to enhance mucus structure integrity during fixation and to initiate polymerization at room temperature for clinical specimen collection. Rheological assessment of Poloxamer 407 polymerization in water (normal formulation) versus formulation in NBF (hereafter, Poloxamer fixative) showed a shift in the polymerization temperature to room temperature (Fig. 2E). In a clinical workflow, Poloxamer fixative is stored at 4°C prior to adding tissue at room temperature (Fig. 2F). The procedure listed was optimized to limit the introduction of extraneous variables and enhance ease of use in the clinic, and we have begun using Poloxamer fixative for limited human colonic biopsy specimen collection at Vanderbilt University Medical Center.

### Poloxamer fixative enables *in situ* analysis of colonic microbes and host cells

To directly compare Carnoy’s solution and Poloxamer fixative for histological applications, a portion of intact colonic tissue from each mouse was divided and fixed separately in each solution. We first tested IF antibodies that performed poorly in Carnoy’s solution, and robust staining was observed with the Poloxamer fixative for PCNA and PCAD (Supp. Fig. 1). To determine which fixation was best for visualizing both the mucus and epithelia, we performed serial multiplex FISH and IF on the same tissue sections using antibodies previously validated in Carnoy’s solution. Five feature multiplex images (GOB5, Eub-eubacterial probe, PCK – pan-cytokeratin, WGA – wheat germ agglutinin, nuclei) are shown for the distal colon, where the highest density of microbes is found (Fig. 3). IF for PCK was used to delineate the epithelial layer (Fig. 3A,B,E,F). GOB5, also known as Calcium-activated chloride channel regulator 1 (CLCA1), is a mucin-processing protein that is secreted with mucus^42^. An intact mucus layer highlighted by antibody staining against GOB5 and WGA was observed with both Carnoy’s solution and Poloxamer fixative (Fig. 3A,C,E,G). Notably, the Eub probe showed greater signal intensity with the Poloxamer fixative compared to Carnoy’s solution (Fig. 3A,D,E,H). Bacteria were only observed with the Carnoy’s solution after increased gain (Fig. 3A - Inset), which exemplifies the challenges with biofilm detection reported with Carnoy’s solution^3,7,43^. Consistent with previous studies^6,8,11^, the Eub signal is absent in the inner mucus layer closest to the epithelium (Fig. 3E,H, white arrow and Supp. Fig. 2). These results show the improved sensitivity for concurrent FISH and IF imaging in colonic tissue with Poloxamer fixative.

**Figure 3:**
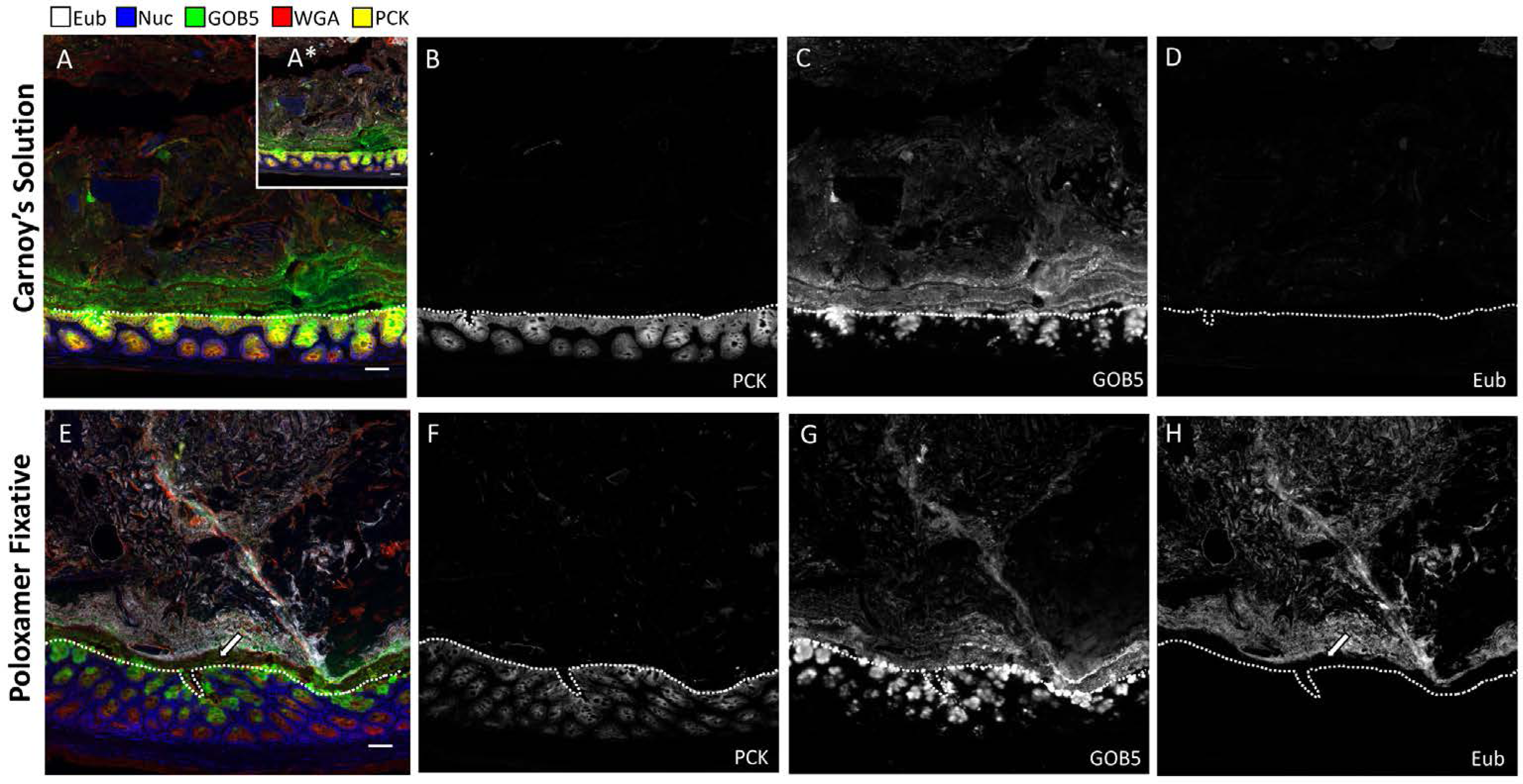
Poloxamer fixation enables concurrent analysis of microbes and tissues, as compared with Carnoy’s fixation. IF of proteins coupled to bacterial FISH over multiple rounds of MxIF, with universal bacterial probe (Eub, white), nuclei (blue), GOB5 (green), lectin (WGA, red), and pan-cytokeratin (PCK, yellow). Selected individual channels (PCK, GOB5, Eub) to visualize epithelium, mucus, and bacteria, with the EUB probe having the same exposure times and gains to compare signal-to-noise in the two conditions. (A-D) Carnoy’s solution, and (E-H) Poloxamer fixative were used for processing the same distal colonic tissue. A* 10X gain was used to detect Eub probe in the Carnoy’s condition. White dotted lines represent epithelial borders. Scale bars = 50 µm.

### Individual microbes are detected in Poloxamer-fixed tissue

To determine the specificity of the Eub probe with Poloxamer fixative, we tested distal colonic samples in parallel with a nonsense probe (Non-Eub). Low intensity staining was observed again with Carnoy’s solution (Fig. 4A), which was further reduced to background level with the non-Eub probe (Fig. 4B). Poloxamer-fixed tissue again had increased Eub staining sensitivity, and at higher magnification, individual, distinct rod-shaped bacteria were observed (Fig. 4C and inset). The non-Eub FISH probe also showed background level of signal in the Poloxamer-fixed tissue (Fig. 4D). These results demonstrate the enhanced sensitivity and specificity of FISH for detection of colonic microbiota with Poloxamer fixation.

**Figure 4:**
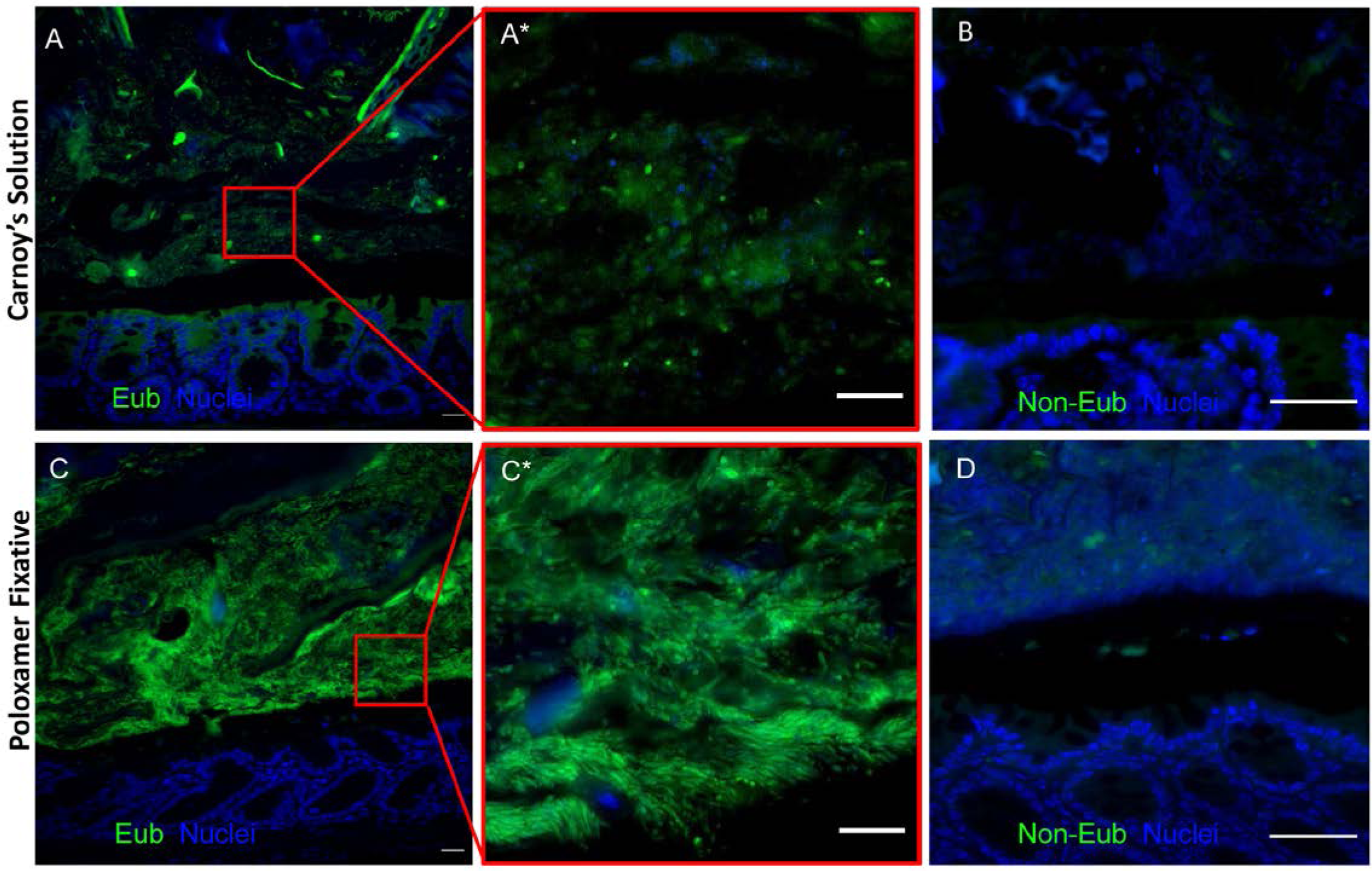
Specific, individual bacteria are identified in Poloxamer-fixed tissue. FISH images of mouse distal colon. (A) Carnoy’s-fixed colon stained with the Eub (green) probe, and (B) non-Eub probe. (C) Poloxamer-fixed colon stained with the Eub (green) probe, and (D) non-Eub probe. A* and C* are higher magnification views of red insets in A and C. Scale bars (A,C) = 20 μm, (A*,B,C*,D) = 10 μm.

### Mucosa-associated biofilms are observed in colonic tissue using Poloxamer fixation

To further validate Poloxamer fixative for biofilm detection with direct implications to the human microbiome, we colonized germ-free (GF) mice with biofilm-positive microbiota slurry isolated from two different human colorectal cancers (Fig. 5A) ^15^. With Poloxamer fixative, bacteria from the first slurry formed dense colonies in direct contact with the colonic epithelium. (Fig. 5B). The second slurry produced a different pattern with bacteria in non-contiguous regions further away from the epithelial cells (Fig. 5C), yet both specimens seemed to lack an inner mucus layer that typically separates luminal bacteria from the epithelium. For our negative control, we observed that no bacteria were detected in sham-inoculated mice, although a mucus layer was maintained (Fig. 5D). More specific FISH probes for *Lachnospiraceae, Bacteroidetes* (CFB), *Fusobacteria*, and *Gammaproteobacteria* demonstrated different abundance patterns in the two inocula (Fig. 5E,F). For example, CFB staining in the first inoculum reflects a relative increase in *Bacteroidetes* (Fig. 5E - inset) compared with the second inoculum (Fig. 5F - inset). *Fusobacteria* were not detected by FISH in any sample, consistent with previous 16S rRNA sequencing at 1 week post-inoculation (Supp. Fig. 3) ^15^. These results demonstrate that Poloxamer fixative enables detection and spatial analysis of mucosa-associated biofilms by multiplex FISH.

**Figure 5:**
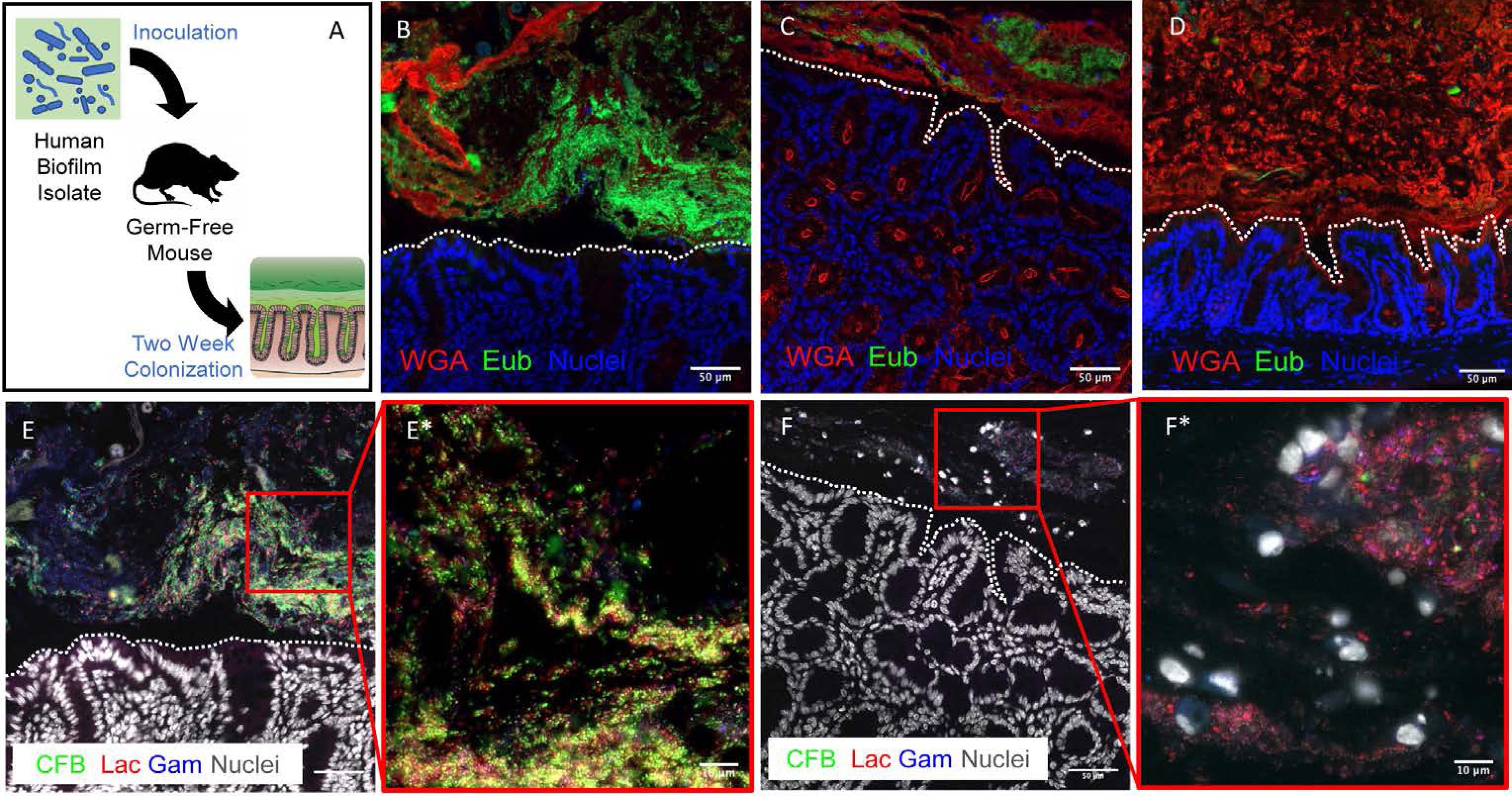
Mouse colons colonized with human CRC patient mucosal homogenate forms mucosa-associated biofilms. (A) A schematic of germ-free (GF) mouse inoculation with human biofilm-positive mucosal slurries. Mice were sacrificed 2 weeks post inoculation and processed using the Poloxamer fixative. Imaging was performed using WGA (red) for mucus, Eub (green) for bacteria, and nuclei (blue) under different conditions of inoculation into GF mice: (B, C) inoculated with two different human isolates, and (D) inoculated with sham treatment. (E and F) Iterative FISH staining rounds with *Bacteroidetes* (green), *Lachnospiraceae* (red), and *Gammaproteobacteria* (blue) probes of the tissue in B and C, respectively. Scale bars = 50 μm. White dotted lines represent epithelial borders. E* and F* are higher magnification views of red insets in E and F. Scale bars = 10 μm.

### Poloxamer fixation permits detection of mucosa-associated biofilms in human colonic adenomas

To demonstrate the clinical utility of our method, portions of human colonic adenomas collected during colonoscopy were immediately placed into Poloxamer fixation for subsequent FISH-based biofilm analysis (Fig. 2F and Supp. Fig. 4). As defined previously^3,12,13,15^, we scored collected patient specimens on a 3-point scale, with a score above 2.5 indicative of biofilm formation (Table 1). Mucosa-associated biofilms were detected on adenomas arising from the ascending colon (Fig. 6). Bacterial invasion into the adenoma was observed in one polyp (Fig. 6C). The advantages of existing repertoires of antibodies and effective bacterial FISH staining enabled robust screening for biofilms in human specimens fixed in the Poloxamer fixative, suggesting that this approach can be a useful clinical tool.

**Table 1:** Biofilm characteristics of human specimens. \***(0 = none, 1 = some, 2 = excellent)** **\*\*(0 = none, 1 =** ≤**5 bacteria, 2 =** ≤**20 bacteria or <200 um long)** **\*\*\*(0-100% of CEC surface)**

| **Sample ID** | **Mucus present\*** | **Bacteria abundance per 200 um CEC\*\*** | **Percentage of biofilm coverage\*\*\*** | **Is there DAPI staining without Cy3?** | **Notes** |
| --- | --- | --- | --- | --- | --- |
| MPP00048A1<br>**HTA11_1429_1011** | 0.5 | 3 | 50% | No | Mostly rods, some cocci in BF; not great mucus though; some autofluorescent blobs (larger than bacteria) to watch out for in some regions |
| MPP00049A1<br>**HTA11_1391_1011** | 1 | 3 | 5% | No | Lots of nice rods everywhere, BF region located within crypts of CEC in one area |

**Figure 6:**
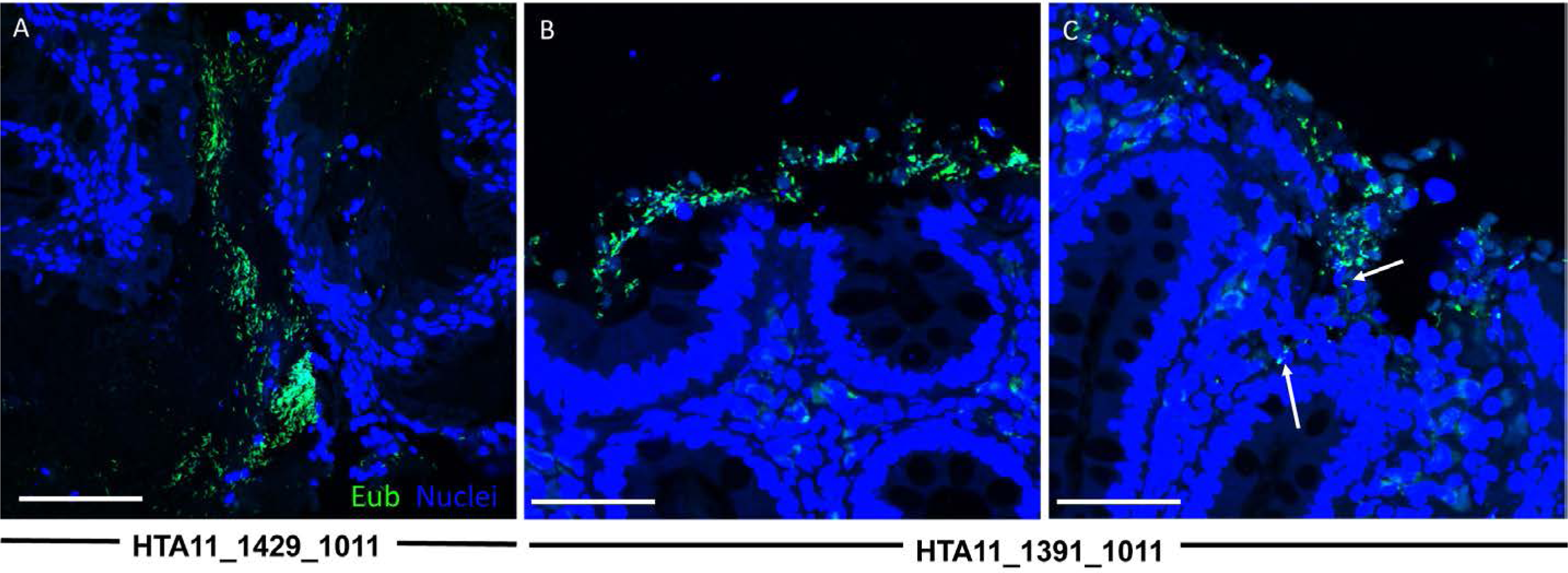
Biofilm positive human colonic polyps demonstrate invasion of bacteria into the mucosa. 63X confocal images of human colorectal polyps fixed in Poloxamer showing regions of biofilm positivity using Eub (green) probes to identify bacteria. (A) Patient sample HTA11_1429_1011. (B and C) Patient sample HTA11_1391_1011. Scale bars = 50 μm.

## Discussion

Microbiota-host interactions are implicated in a range of disorders, including CRC. Specifically, human CRC-associated stool microbiota transplanted into GF or carcinogen-treated mice exacerbates colon epithelial cell proliferation and the tumor-associated immune response^44^. Colonic mucosa-associated biofilms are defined by the direct interaction between bacteria and epithelial cells. They are present in approximately 89% and 12% of right-sided and left-sided colon tumors, respectively^3^. CRC in human familial adenomatous polyposis appears to be uniformly biofilm-positive^13^. Human colonic biofilms are carcinogenic in GF ApcMin mice^15^. With a reported range of 13–35% colonic biofilm positive in healthy subjects, these findings indicate clinical relevance for using biofilm to stratify high-risk pre-cancer lesions.

The established tissue fixation for detecting colonic biofilms is Carnoy’s solution. A major limitation of Carnoy’s solution is that IF sensitivity is reduced and inadequate for detecting epithelial antigens (Fig. 1). Generally, dehydration fixation approaches are able to retain mucus in samples, but they also reduce the efficacy of FISH staining^11,45^. Conventional FFPE tissue is limited by the poor preservation of the mucus layer (Fig. 1). Our goal was to develop a broadly applicable reagent that preserves mucus and biofilms without compromising IF detection of host factors. With the Poloxamer fixative, we overcame the need to apply heat for polymerization and developed a clinically compatible, room temperature reagent (Fig. 2).

We demonstrated how Poloxamer-fixed samples were superior to Carnoy’s-fixed samples for visualizing both mucosa-associated bacteria and host cells in the same tissue section (Fig. 3). Whereas agarose has been shown to also provide physical scaffold^36^, the physical and chemical properties of Poloxamer make it the ideal substrate for preservation of mucus. The Poloxamer fixative polymerizes at room temperature, enabling its incorporation into existing clinical protocols for human studies. Methods that enable multi-modal analysis will become very valuable in future endeavors for understanding complex tissue ecosystems. Our efforts represent an advance in the field of multiplex imaging, and one of the first to simultaneously evaluate composition and organization of both microbes and host cells

Previous assessments of biofilms relied on consensus of experts to score samples for determining biofilm positivity^3,12,13^. The increased FISH signal observed with Poloxamer fixative potentially enables the identification of biofilms with improved technical proficiency and machine learning. Unlike sequencing approaches to characterize the microbiome, FISH requires pre-selection of candidates, but it provides spatial information important for determining microbial behaviors^7,14^.

To our knowledge, this is the first use of Poloxamer 407 for tissue fixation^37,39,40^. Our successful implementation will enable its wider deployment in clinical pipelines for human microbiome and biofilm research.

## Author Contributions

MCM conceived the study, performed experiments, analyzed the data, compiled the figures, and wrote the manuscript. JLD performed microbiome transplant experiments, scored human specimens, and contributed to the writing of the manuscript. NOM assisted with mouse experiments, intellectually contributed to the study, and wrote the manuscript. AJS contributed to the study design, and assisted with experiments. JTR assisted with imaging. PNV and CRS assisted with mouse experiments. RJC contributed to funding and writing of the manuscript. MJS facilitated human specimen collection, and contributed to funding and writing of the manuscript. CLS supervised the research, contributed to the study design and writing of the manuscript. KSL conceived the study, analyzed the data, wrote the manuscript, obtained funding, and supervised the research.

## Acknowledgements

The authors would like to thank Dr. Cynthia Reinhart-King for advice on polymers, Katarzyna Zienkiewicz and Dr. Scott Guelcher for rheometer use, and Dr. Eliot McKinley for his advice on MxIF use. KSL and AJS are funded by R01DK103831. RJC is funded by P50CA236733 and R35CA1975703. MCM, JLD, JTR, RJC, MJS, CLS, and KSL are funded by U2CCA233291. NOM is funded by T32DK007673. PNV is funded by T32HD007502. Participant recruitment and biospecimen collection were conducted in part by the Survey and Biospecimen Shared Resource which is supported in part by P30CA68485. Tissue fixation and processing was provided by the NCI Cooperative Human Tissue Network (CHTN) Western Division UM1CA183727.

## Materials and Methods

### Tissue Fixation and Processing

For Carnoy’s fixation, tissues were immersed in Carnoy’s solution (6:3:1 ratio of ethanol:glacial acetic acid:chloroform) and processed using standard procedure^6^. Tissues were also fixed using standard NBF fixation. For Poloxamer fixation, the Poloxamer 407 solution was made at 4°C, where 20% w/w Poloxamer was mixed into 10% NBF. Aliquots were kept on ice prior to the addition of the tissue specimen. Once the specimen was immersed in the solution, the mixture was brought to room temperature at 25°C and allowed to polymerize. The mixture was visually inspected for polymerization by checking for the lack of movement when the conical tube was inverted. From here, the tissue was fixed at room temperature for 24-hour, followed by standard histological processing. Tissue blocks prepared from fixed tissues were sectioned at 5 microns onto slides ^22^.

### Rheological Testing

To test for polymer gelation properties, rheological testing was performed using Rheometer AR2000 ex machine with both the Poloxamer fixative formulation (20% w/w Poloxamer 407 into 10% NBF) and normal Poloxamer formulation (20% w/w Poloxamer 407 into water). To test sol-gel transitions, temperature was modulated from 5°C to 40°C with controlled stress and rate^38,40^, and half gelation temperature was calculated by a dose response non-linear log fit in Prism (Graphpad).

### Tissue Collection

Conventional mouse experiments were performed under protocols approved by the Vanderbilt University Animal Care and Use Committee and in accordance with NIH guidelines (Protocol: M1600047). Animals were euthanized and colonic tissues were collected using previously published procedures^46,47^. Germ-free mouse experiments were performed at Johns Hopkins University under protocols approved by the Johns Hopkins University Animal Care and Use Committee (Protocol: MO17M111) in accordance with NIH guidelines.

Human colonic tissues were collected from participants in the Colorectal Molecular Atlas Project (COLON MAP), a study approved by Vanderbilt Institutional Review Board (Protocol: 182138). All participants provided written informed consent. Participants underwent colonoscopy as part of their routine care following standard bowel cleansing preparation. Polyps were removed according to standard of care and bisected. Visually normal mucosal biopsies were obtained in the mid-ascending colon using jumbo biopsy forceps. To increase likelihood of biofilm positivity, only participants with a polyp in the cecum or ascending colon were included in this analysis. Each tissue (either normal biopsy or polyp portion) was placed in individual labelled tubes of Poloxamer fixative, and gently inverted twice to ensure full coverage of the sample.

### Multiplex fluorescence imaging

Sequential antibody staining and dye inactivation was performed as described^22^. FISH probes and antibodies used are listed in **Tables 2 and 3**. Modifications to the protocol includes the incorporation of FISH, as described below, prior to immunofluorescence. Fluorescence of FISH probes was inactivated using the same methods as antibodies. For comparing Carnoy’s fixation and Poloxamer fixation, tissues were processed simultaneously and imaged with the same exposure time with the Olympus X81 inverted microscope with a motorized stage and with filter sets specific for DAPI, GFP, CY3, CY5, and CY7. Images were also collected on Zeiss Axio-Imager M2 microscope.

**Table 2:** Antibodies used to detect host cells.

| **Marker** | **Channel** | **Exposure** | **Concentration** | **Product ID** | **Clone/Cat #** | **Clonality** | **Host Species** |
| --- | --- | --- | --- | --- | --- | --- | --- |
| PCK | FITC | 100 ms | 1:200 | ab11214 | PCK-26 | Monoclonal | Mouse |
| GOB5 | 647 | 100 ms | 1:200 | ab180851 | EPR12254-88 | Monoclonal | Rabbit |
| PCNA | 488 | 200 ms | 1:100 | CS 8580S | P10 | Monoclonal | Mouse |
| PCAD | Cy3 | 100 ms | 1:100 | DC13090 | RB-9036 | Polyclonal | Rabbit |

**Table 3:**
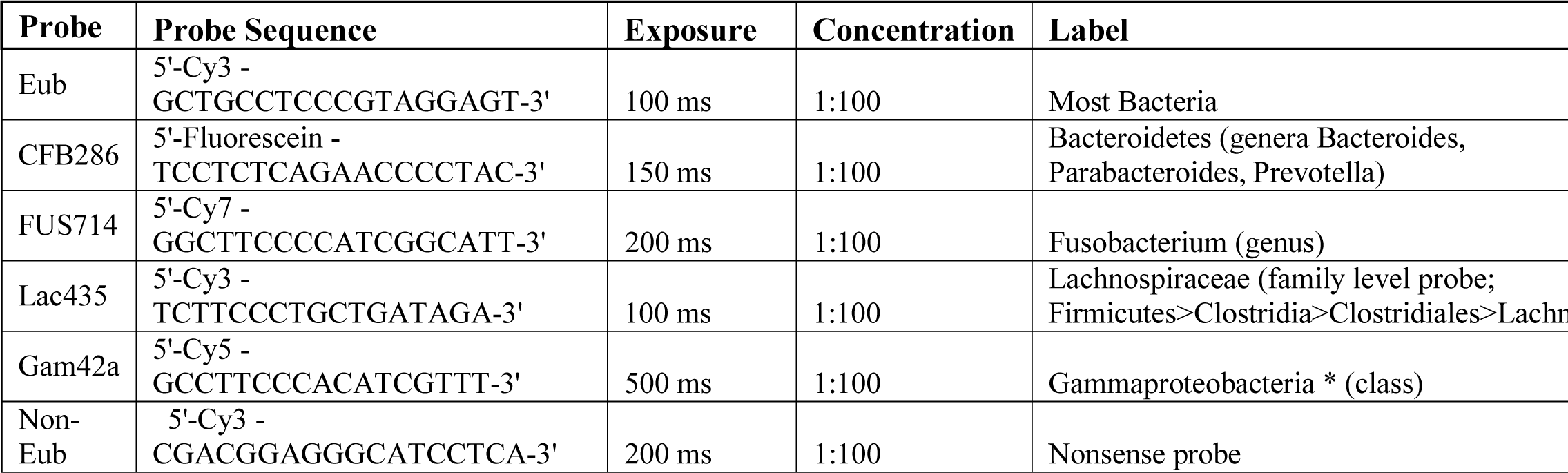
Nucleic acid probes used to microbes.

### Fluorescent *in situ* hybridization

FISH probes were suspended in sterile water at a concentration of 0.2 nmol/ul (200 uM) and stored at −20°C. Hybridization buffer (20 mM Tris-HCl [pH 8.0], 0.9 M NaCl, 0.01% sodium dodecyl sulfate) and FISH wash buffer (225mM NaCl, 20mM Tris, 5mM EDTA) were pre-warmed in a 46°C oven. Slides were then de-paraffinized through 3 × 5 min dips in Histoclear under the fume hood. Re-hydration of slides was done using an ethanol gradient. Slides were placed for 5 minutes in each of the following solutions: 100% ETOH, 100% ETOH, 95% ETOH, 95% ETOH, 70% ETOH, followed by storage in 20 mM Tris buffer. Probes were diluted to 2 μM in hybridization buffer prior to application. Approximately 100 ul volume was applied per sample, to fully cover the fixed tissue. Slides were then incubated for approximately 1.5-2 hours at 46°C in a humidified chamber (1.5 hours for the universal probe; 2 hours for specific taxa probes). Slides were washed with FISH wash buffer 3 × 5 minutes on shaker.

### Biofilm Screening

Patient tissues were collected and screened for biofilms using the universal bacterial probe (EUB338), as described previously^3,12,13^. Briefly, tissues were defined as biofilm positive if there were more than 10^9^ bacteria/ml invading the mucus layer (within 1 μm of the epithelium) for at least 200 μm of the epithelial surface.

### Human tissue inocula preparation

The study design was developed to give priority to reproducibly test the hypothesis that human mucosal microbiota populations from different patients will display unique mucosal interactions in a mouse model. Thus, to allow for limitations in available human tissues and GF mice, we used inocula composed of tissue from 2 patients. Inocula were prepared from 3 mm diameter tissue pieces that were collected from the resected CRCs of the two patients, snap-frozen, and stored at −80°C. All inocula were prepared anaerobically by mincing and homogenizing tissue pieces in PBS in an anaerobic hood to a dilution of 1:20 (weight/volume). Prior to gavage into germ-free mice, the inocula were further diluted 1:10 in PBS, for a final of 1:200 w/v dilution from the original tissue.

### Mouse colonization

Germ-free (GF) C57BL/6 wild type animals were transferred to gnotobiotic isolators (separate isolator for each experimental group) and gavaged with 100 μl of human tissue slurry inoculum. Mice were euthanized at indicated time points^15^. About 1 to 2 × 0.5 cm snips were taken from the proximal and distal colon, and then fixed in accordance to the protocols above. Tissue was processed, paraffin-embedded, and sectioned.

### Data availability statement

All data generated or analyzed during this study are included in this published article (and its supplementary information files).

## Supplementary Figure Captions

**Supplementary Figure 1: Poloxamer fixative preserves antibody staining** Comparison between Carnoy’s and Poloxamer fixation of (A) PCNA, (B) pan-Cadherin (PCAD). Scale bars = 50 µm.

**Supplementary Figure 2: Closer examination of Poloxamer fixed mouse colon reveals intact inner mucus layer devoid of microbes**.

Mouse colon fixed using the Poloxamer fixative and stained with GOB5 (red) and Eub (green). Black arrow points to intact mucus layer. Scale bars = 50 μm and 20 μm (inset).

**Supplementary Figure 3: Lack of Fusobacterial staining observed in mouse colon colonized with human CRC patient mucosal homogenate**. Iterative round of FISH staining for *Fusobacteria* (green) probe of the tissue in Fig.5B. Scale bars=50 μm and 10 μm (inset).

**Supplementary Figure 4: Normal human biopsy in fixed Poloxamer fixative reveals preserved architecture of the colonic mucosa**. Scan of a human colon biopsy stained using WGA (red) for mucus and Eub (green) for bacteria. Scale bar = 50 µm.

