## Supplementary figures and images for "Clinically adaptable polymer enables simultaneous spatial analysis of colonic tissues and biofilms"

### Supplemental Figures

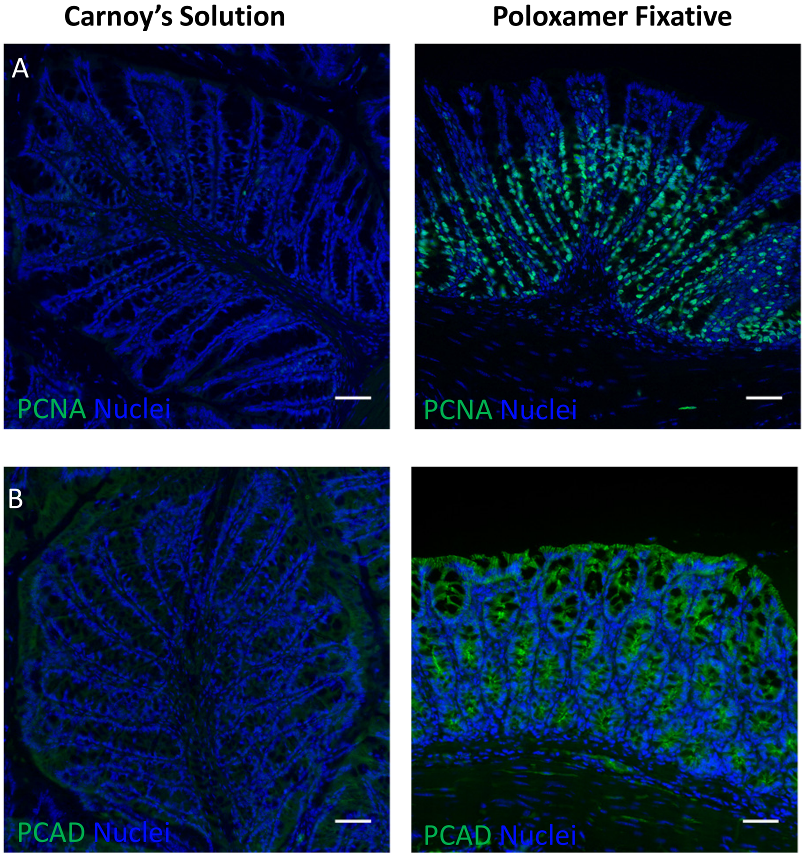


Supplementary Figure 1

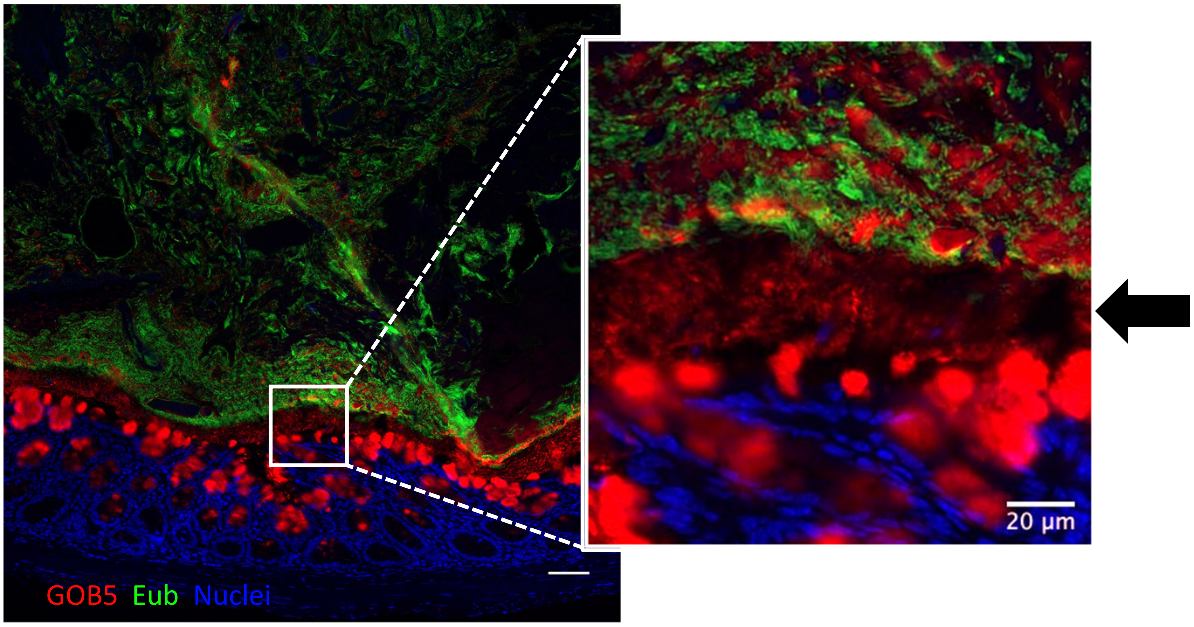


Supplementary Figure 2

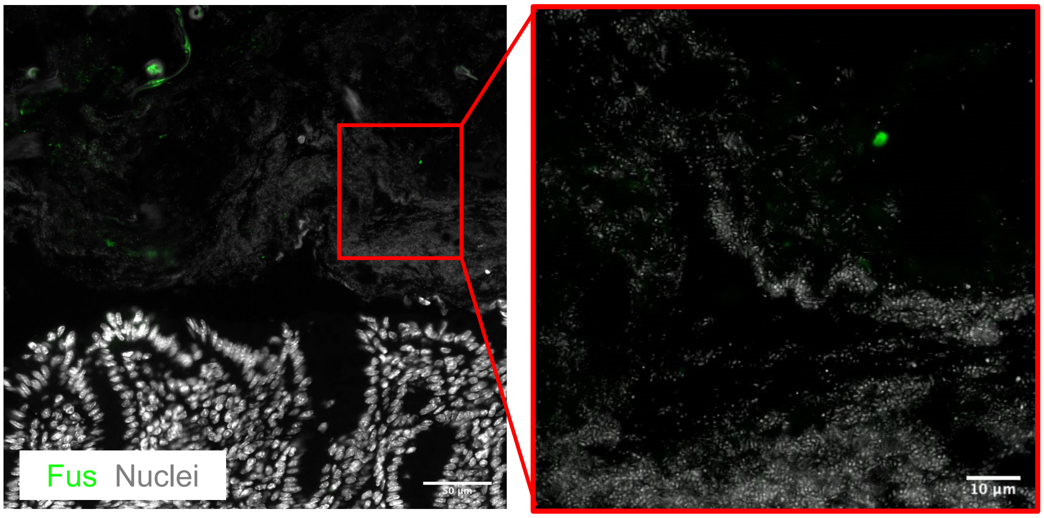


Supplementary Figure 3

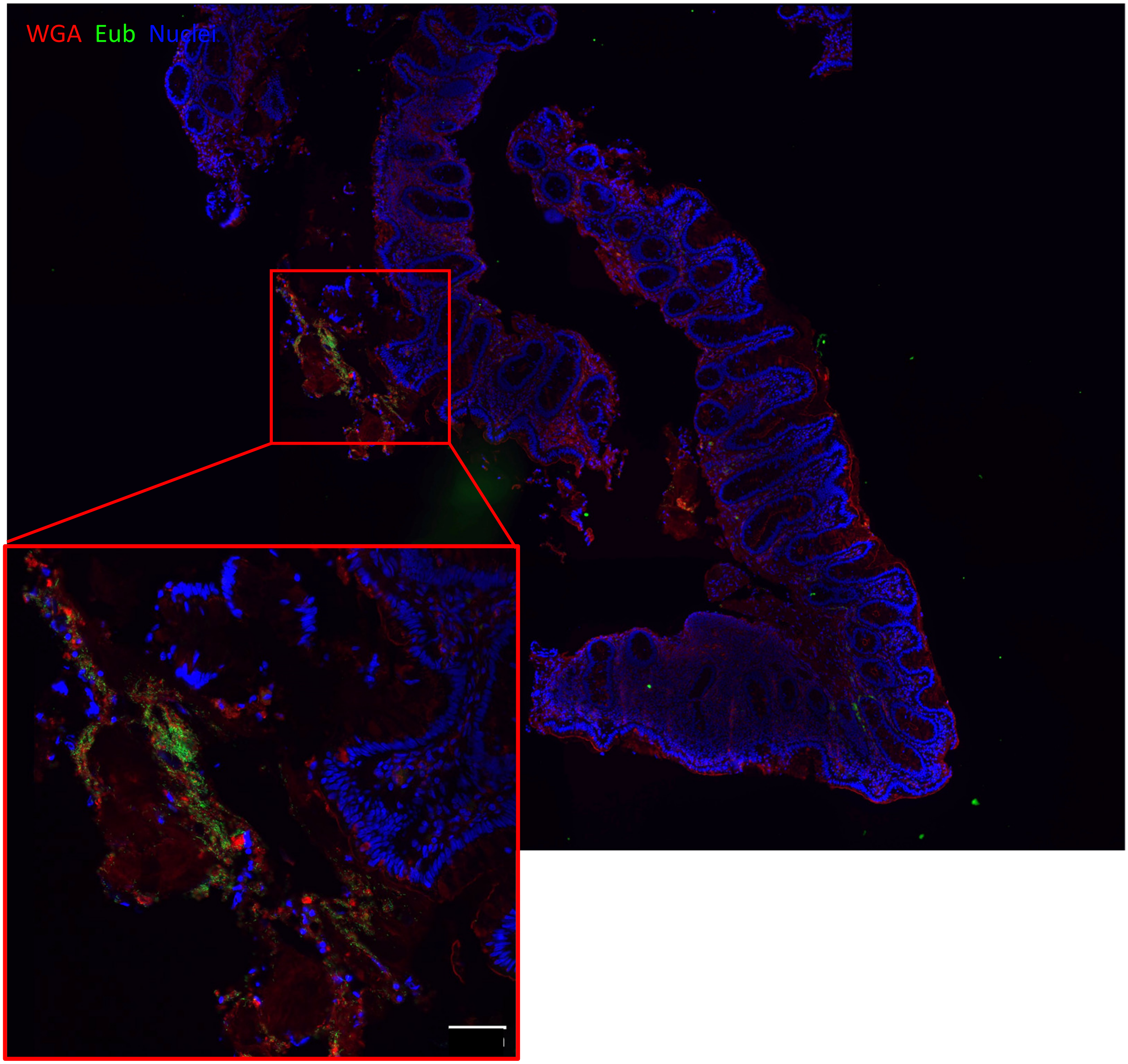


Supplementary Figure 4
